# Tooth Shape Adaptations in Aglyphous Colubrid Snakes Inferred from 3D Geometric Morphometrics and Finite Element Analysis

**DOI:** 10.1101/765719

**Authors:** Mahdi Rajabizadeh, Sam Van Wassenbergh, Christophe Mallet, Martin Rücklin, Anthony Herrel

## Abstract

To date there are few detailed and quantitative studies investigating the evolution of the tooth shape and function in Aglyphous snakes in relation to diet. To study dental adaptations to diet, a lineage that is of particular interest due to its large range of adult body sizes, is the one including dwarfed snakes of the genus *Eirenis* and their immediate sister group, whip snakes of the genus *Dolichophis.* A considerable evolutionary decrease in the size is observed from a *Dolichophis*-like ancestor to the miniature *Eirenis*, coupled with a considerable shift in their diet from a regime consisting mainly of endotherms with endoskeleton to ectotherms bearing a hard exoskeleton. Maxilla, palatine, pterygoid and dentary teeth were examined in an adult and a juvenile of *Dolichophis schmidti*, one *Eirenis punctolineatus* and one *Eirenis persicus*. 3D Geometric Morphometrics comparison revealed maxilla and palatine teeth of the *E. persicus* are blunt and conical shape while those teeth are sharp and elongated in *E. punctatolineatus* as well as the adult and juvenile *D. schmidti*. A similar difference could be noted for the pterygoid teeth. In contrast, the dentary teeth are not as different among the examined snakes. Blunt and conically shaped teeth, as observed in *E. persicus*, seem to be more adapted for biting hard bodied, arthropod prey, while sharp and elongated teeth in *Dolichophis* and *E. punctatolineatus*, are specialized for puncturing endotherm prey. The results of a finite element analysis confirms that during biting a hard bodied prey, the generated stresses in *E. persicus* tooth is mostly confined to the tip of the tooth and mostly well below the von Mises yield criterion the tooth. In contrary, *D. schmidti* tooth appears less well suited for biting a hard prey since the generated stresses widely distribute across the tooth with values roughly 2 to 3 times higher than the von Mises yield criterion of the tooth. A lower degree of specialization that was observed among the dentary teeth in the examined snakes suggest a similar functional constraint in pushing the prey against the upper tooth rows.

## Introduction

Vertebrate teeth refer to highly mineralized appendages in mouth, mainly associated with ingestion and processing of the prey, but they also frequently serve other functions, such as defense(Koussoulakou et al., 2009). Phylogeny and dietary habits have driven the teeth of vertebrates to acquire numerous anatomical forms and shapes (Knox and Jackson, 2010). Snake teeth are recurved, sharply pointed, and mainly acrodont, although some are pleurodont, as in scolecophidians (Zaher and Rieppel, 1999). Snakes can be placed in four groups based on their dental morphology. Aglyphous snakes have a series of more or less similar backwardly curved teeth on the maxillae, without a groove. Opisthoglyphous snakes possess a full row of teeth on the maxillae, including enlarged, grooved (on the mesial or lateral tooth surface), backwardly curved teeth in the middle of the row or at the back. Proteroglyphous snakes have fixed, enlarged teeth on the anterior portion of the maxillae with venom grooves that except for an opening near the tip, with the rest of the groove being closed. Solenoglyphous snakes that have tubular, enlarged, fully closed teeth on the reduced maxillae that can be rotated around the prefrontals (Berkovitz and Shellis, 2016; Deufel and Cundall, 2006; Kardong and Young, 1996). In Aglyphous snakes, which comprise roughly 85% of extant snakes, teeth are considered as not specialized for venom injection nor to mediate venom penetration into the prey body (Berkovitz and Shellis, 2016).

Previous studies have reported structural and functional adaptations in Aglyphous snakes in relation to diet, including the occurrence of long, numerous, and very sharp teeth in some piscivore snakes (Savitzky, 1983), the occurrence of hinged, weakly ankylosed teeth in some lizard (Savitzky, 1981; Savitzky, 1983) or arthropod eating snakes (Jackson et al., 1999), the occurrence of reduced teeth in the size and number in egg-eating snakes (Gans, 1952) or elongated and numerous dentary teeth as well as reduced palatomaxillary teeth in snail-eating (cochleophages) snakes (Savitzky, 1983). To date there are few detailed and quantitative studies investigating the evolution of the tooth shape and function in Aglyphous snakes in relation to diet.

A lineage that is of particular interest due to its large range of adult body sizes, is the one including dwarfed snakes of the genus *Eirenis* and their immediate sister group, whip snakes of the genus *Dolichophis* (Rajabizadeh et al., 2019). *Dolichophis* snakes (maximum size: about 2500 mm) feed on a variety of food items, including small mammals, birds, lizards, and more rarely on bird eggs, arthropods and even other snakes (Göçmen et al., 2008; Lelièvre et al., 2012; Rajabizadeh, 2018; Terent év and Chernov, 1965). *Eirenis* snakes are classified in four subgenera, *Eirenis*, *Pediophis*, *Pseudocyclophis* and *Eoseirenis*, which except for the latter subgenus, are more or less dwarfed and feed on lizards and/or arthropods (Rajabizadeh, 2018). *Eirenis* (*Pediophis*) *punctatolineatus* (maximum size: 758 mm) feeds mainly on lizards and arthropods, while *Eirenis* (*Pseudocyclophis*) *persicus* (maximum size: 371 mm) feeds nearly exclusively on arthropods (Rajabizadeh, 2018; Terent év and Chernov, 1965).

Hence, a considerable evolutionary decrease in the size is observed from a *Dolichophis*-like ancestor to the miniature *Eirenis*, coupled with a considerable shift in their diet from a regime consisting mainly of endotherms with endoskeleton to ectotherms bearing a hard exoskeleton. We hypothesized that 1) tooth shape in miniature *Eirenis* snakes is different than in their *Dolichophis* like ancestors; 2) *Eirenis* tooth shape reflects a suite of structural and functional adaptations to better resist the strains associated with the biting of harder prey than *Dolichophis*. To test these hypotheses, we provide a detailed structural and functional comparison of the shape of the teeth in *Dolichophis* and *Eirenis*.

## Materials and Methods

### Specimens

To study dental adaptations to diet we examined an adult and a juvenile *D. schmidti*, one *E. punctolineatus* and one *E. persicus*, all of which were micro CT-scanned. Biometric data of the examined specimens are presented in Table 1.

**Table 1.** Biometric data as well as micro-CT scan details of the examined specimens. Acronyms are as follows: snout-vent length (SVL), total length (TL) and head length (HL).

| Species | SVL<br>(mm)* | TL<br>(mm) | HL<br>(mm) | Nr. of Projections | Voxel Size<br>(μm) |
| --- | --- | --- | --- | --- | --- |
| *Dolichophis schmidtii*<br>(adult) | 750 | 150 | 35.2 | 1762 | 19.863 |
| *Eirenis punctatolineatus* | 408 | 136 | 14.8 | 1781 | 11.261 |
| *Eirenis persicus* | 233 | 66 | 8.0 | 1802 | 6.2 |
| *Dolichophis schmidtii*<br>(juvenile) | 380 | 92 | 12 | 1714 | 7.616 |

\*: Millimeter

The micro-CT scans of the heads of four snake specimens were performed at the Centre for X-ray Tomography of Ghent University (Masschaele et al., 2007). The setup was a transmission head of a dual-head X-ray tube (Feinfocus FXE160.51) and a-Siflat panel detector (PerkinElmer XRD 1620 CN3 CS). The focal spot size was 900 nm at a tube voltage of 130 kV for high resolution. Number of projections and voxel size of the scanned specimen is presented in Table 1. Exposure time was 2 seconds per projection, resulting in a 360° output CT Scan. The raw data were processed and reconstructed using the in-house CT software Octopus (http://www.octopusreconstruction.com; Vlassenbroeck et al., 2007) and rendered using Amira V. 5.4.1 (Mercury Systems of Visage Imaging GmbH). The CT-rendered images were color coded to distinguish separate ossified units, where stiff and rigidly interconnected bones were given a single color. CT scan details of the scanned specimens are presented in Table 1.

The palatomaxillary arch and the mandible of each examined specimen were isolated from the rest of skull using the segmentation tool in AMIRA. Teeth of the left side of the palatomaxillary arch and the left mandible were isolated along a straight line crossing the anterior and posterior edge of the tooth’s socket and then prepared using Geomagic Wrap V. 2017.1. To prepare a tooth for 3D shape analysis, the basal edges of each tooth were covered by a flat plane, except for the central pulp cavity. Then, each tooth was checked for holes, spikes, self-intersection and non-manifold edges using the Repair Module in Geomagic. The surface noise due to scanning was reduced (mid smoothness level) using the Smooth Module in Geomagic Wrap. Since snake teeth are replaced regularly, only those active teeth, fully fused into the socket, were used in this research.

### Morphometrics

A combination of the anatomic landmarks and semi-landmarks sliding on both curves and surfaces (Gunz and Mitteroecker, 2013; Gunz et al., 2005) were used to quantify the shape variation of the teeth. A total of 306 landmarks were placed on each tooth including 10 anatomical landmarks and 296 sliding semi-landmarks. Of those 120 were curve semi-landmarks bordering the base of the tooth, and 176 were surface semi-landmarks distributed homogenously over the tooth surface (Figure 1).

**Figure 1.**
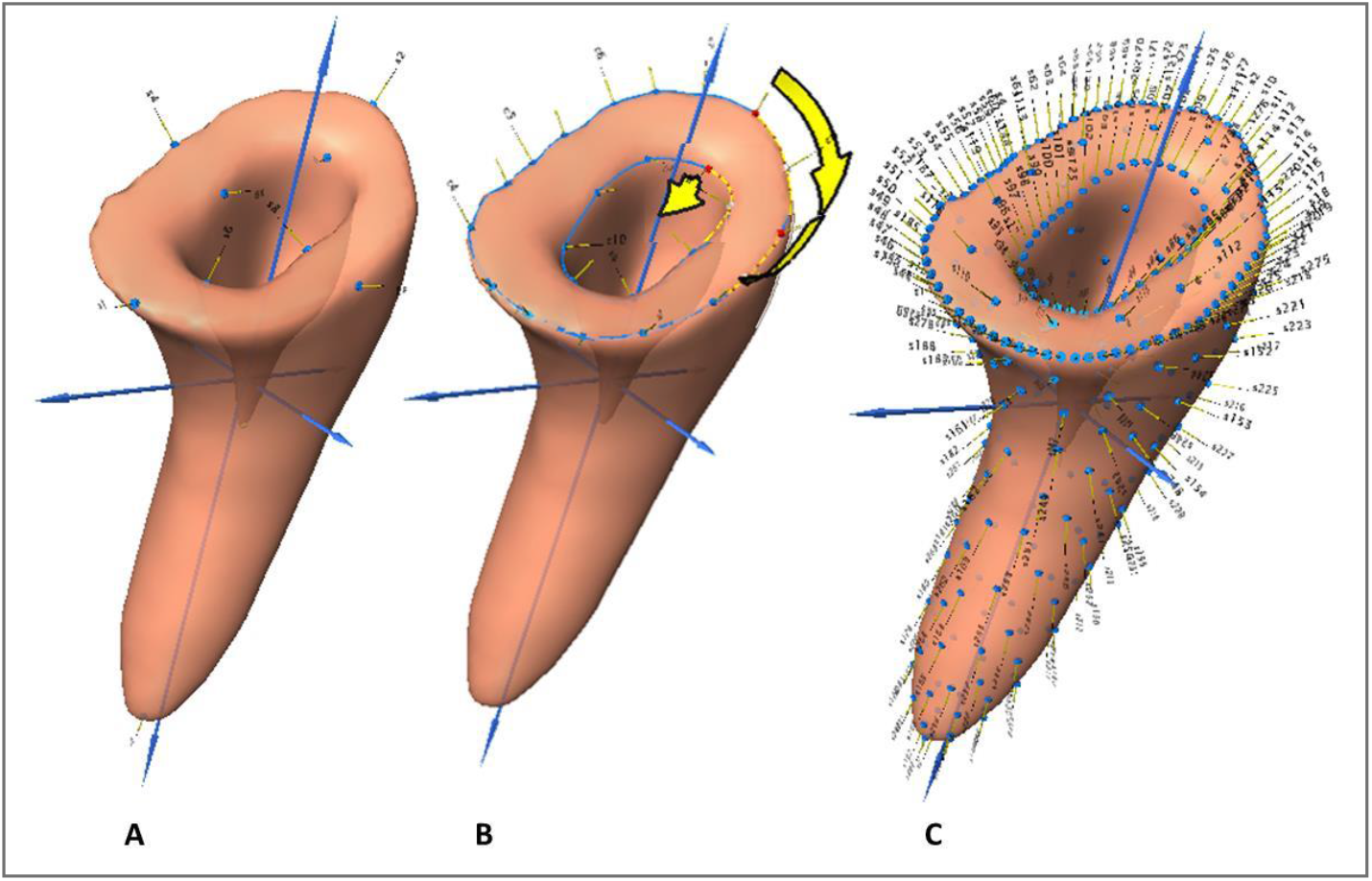
Position of the landmarks on a tooth, including 10 anatomical landmarks (A); 120 curve semi-landmarks (eight curves on outer edge of the tooth base, specifying 80 curve semi-landmarks, and four curves on inner edge of the tooth pulp specifying 40 curve semi-landmarks) (B); 176 point semi-landmarks on the tooth surface.

The anatomic landmarks and semi-landmarks were digitized using the IDAV Landmark software package (Wiley et al., 2005) for each tooth. Curve sliding semi-landmarks were equidistantly re-sampled and bordered by anatomical landmarks (Botton-Divet et al., 2016; Gunz et al., 2005). Surface semi-landmarks were digitized manually on the surface of a template tooth mesh using the IDAV Landmark software package. This template was used to projection surface semi-landmarks onto the surface of all the examined teeth. As recommended by Gunz and Mitteroecker (2013), curve and surface semi-landmarks were slid to ensure that each point can be considered as geometrically homologous. Sliding was done minimizing the bending energy of a Thin Plate Spline between each specimen and the template at first and then the Procrustes consensus was used as a reference during the next three iterative steps. Sliding of surface semi-landmarks over the surface of the teeth was performed using the “Morpho” package (Schlager, 2013) as detailed in Botton-Divet et al. (2015). Both curve and surface semi-landmarks were slid using the bending energy minimisation criterion.

A generalised procrustes analysis was performed using the “Morpho” R package(Schlager, 2013). Shape variation was visualized using a Principal Component Analysis (PCA), performed on the superimposed coordinates projected onto the tangent space, using “Morpho” R package(Schlager, 2013). The analyses were performed in R (Team, 2018).

As a proxy of shape divergence among the teeth of the examined snakes, Euclidean Distances along the centroid PC scores were computed. The average of PC scores on the first and second PC were extracted. Subsequently the distance between the PC scores was computed using the following formula:

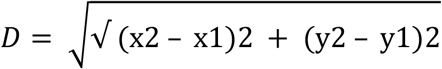

### Finite Element Analysis (FEA)

To evaluate the effect of tooth shape in resisting the deformations imposed during biting a hard prey, a Finite Element Analysis was performed. In this analysis we focused on maxillary teeth, since, 1) a clear difference in the shape of maxillary teeth of the examined specimens was documented, 2) some data are available to evaluate the movement of the maxillary bone during the biting of a prey (Cundall, 1983; Schwenk, 2000), 3) muscular structures involved in maxillary biting are rather straightforward simplifying the orientation of the external forces acting on the maxilla.

### Geometry preparation

The geometric mean shape of the maxillary teeth of *D. schmidti* and *E. persicus* were selected based on the PC1 vs. PC2 scatter plot (Figure 4). Hence, 8^th^ maxillary teeth of *E. persicus* and 5^th^ maxillary teeth of *D. schmidti* (both on the left maxilla) were selected as the maxillary tooth mean shape. The mean shape was trimmed using Fix Module in 3-Matic Medical V. 11 (Materialise, Leuven, Belgium).

### Meshing

To calculate the proper mesh size, a convergence analysis was performed on the maxillary tooth mean shape of *E. persicus*. For this analysis, 0.00024 N was implemented along the main axis of the tooth, over three outermost nodes on the tip of the tooth. Constraint and loading conditions are explained in a following section. The resulting plot shows the element size against the maximum computed stress (von Mises) and confirmed that the variation around the element size equal to 0.01 millimeter does not affect the resulting stress values drastically Figure 2). Using the element size 0.01 mm, the maxillary tooth mean shape of *E. persicus* was meshed with 7176 tetrahedral elements and the maxillary tooth mean shape of *D. schmidti* was meshed with 13052 tetrahedral elements. Both the meshes generated with the default growth rate of 1.2, using Meshing Module of ANSYS v 15.0 0 (ANSYS, Canonburg, PA, USA).

**Figure 2.**
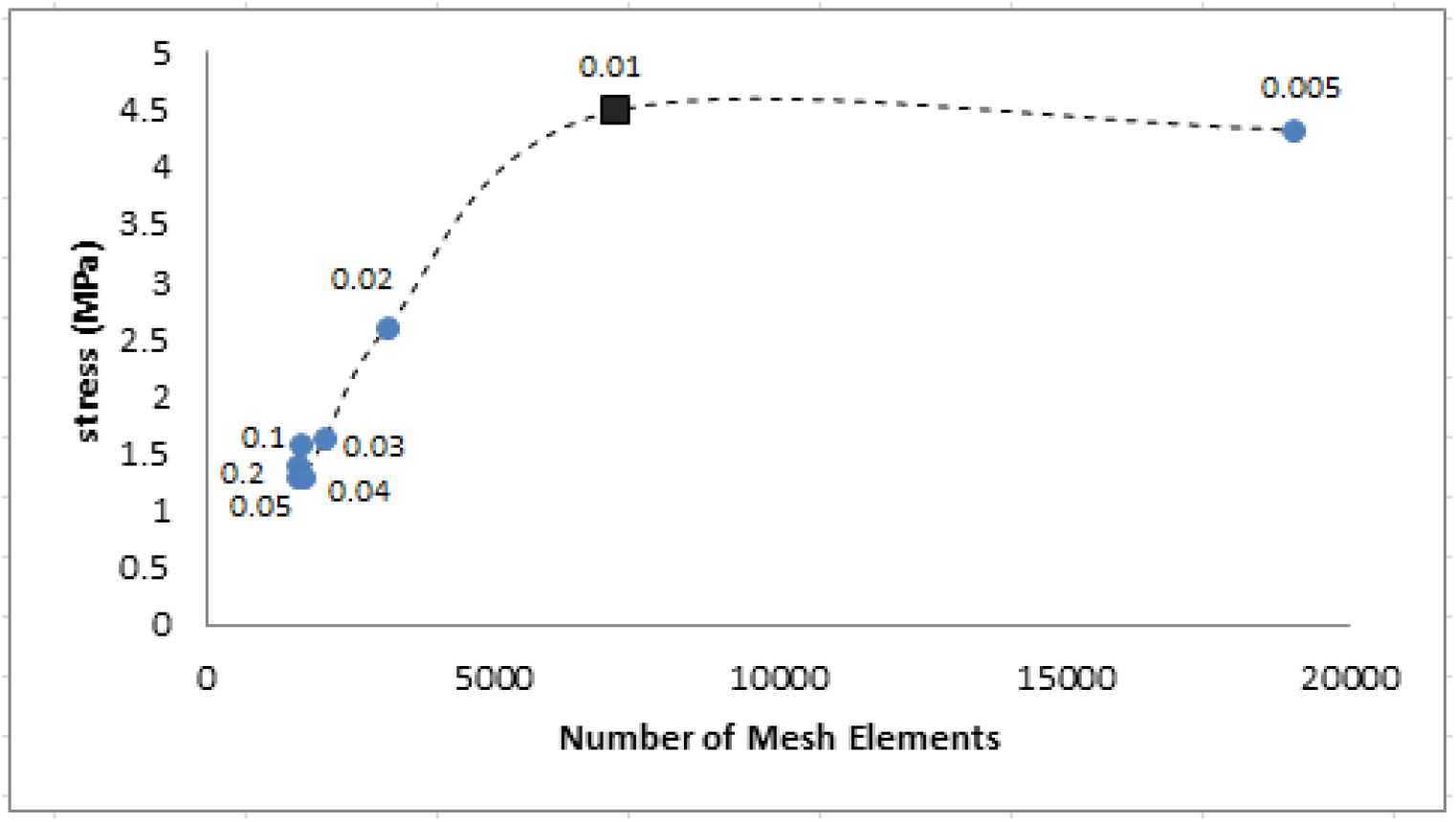
Mesh convergence analysis on the maxillary tooth mean shape of *E. persicus* plotting the element number against the maximum computed stress (von Mises). Respected element size is noted over the points. The square represents the element size equal to 0.01 mm.

### Material properties

Despite the extensive data about the material and mechanical properties of human teeth (Zhang et al., 2014), little is available about the material properties of teeth in other vertebrates. Teeth have been shown to be anisotropic (Waters, 1980) and a complex structure of enamel and dentine layers has been reported in reptiles (Zahradnicek et al., 2014). However, data concerning the thickness and distribution of enameloid and dentine are not available for the examined colubrid snakes. Hence, because of comparative nature of our model, we considered the examined teeth as static, linearly elastic, isotropic and homogenous, for the sake of model simplicity. There are no data summarizing Young’s modulus and Poisson’s ratio for reptile teeth. Hence, we used the data available for human dentine: Young’s modulus (E=18000MPa) and Poisson’s ratio (v=0.31) (Bessone et al., 2014).

### Constraint, forces and loading condition

The application of the realistic forces and constraints are essential for successful FEA modeling (Dumont et al., 2005). In the *Eirenis* and *Dolichophis* snakes, prey ingestion is mainly performed by the maxilla. Each maxilla is a curved bone, posteriorly connected to the ectopterygoid and medially articulated with the ventral surface of the prefrontal. Hence, the maxilla is a movable bone, with a degree of rotation along the medial articulation with the prefrontal. Maxilla movement comes from the contraction of pterygoideus and pterygoideus accessories muscles (Kardong et al., 1986). During prey capture the maxillary teeth slide over the prey to align forces acting on the teeth with the long axes of the teeth (Schwenk, 2000). Hence in the movable maxilla, a range of the direction of the forces, acting around the main axis of a maxillary tooth can be expected. Given the above-mentioned condition, we set up the model as following:

#### Constraint

The external nodes on the tooth base were designated as fixed to constrain both the translational and rotational displacement.

#### Force

In *Eirenis persicus*, maximal force generated by pterygoidous and pterygoideus accessories muscles was calculated as 0.19 N (Rajabizadeh et al., 2019). Since in the current study, the left maxillary bone of *E. persicus* has eight active teeth, firmly attached to the maxilla, and assuming that all teeth are in contact with the food item during the ingestion, a force of was applied to each tooth.

#### Loading condition

The force was distributed over the outer surface of the tip of the tooth. Considering the geometry and sharpness of the teeth, the force was implemented over 75 nodes on tip of the *E. persicus* tooth and 31 nodes on tip of the *D. schmidti* tooth. The direction of the force was set up along the sagittal surface of the tooth, across the main axis of the tooth, as well as ±45° off the main axis, at 15° intervals (Figure 3).

**Figure 3.**
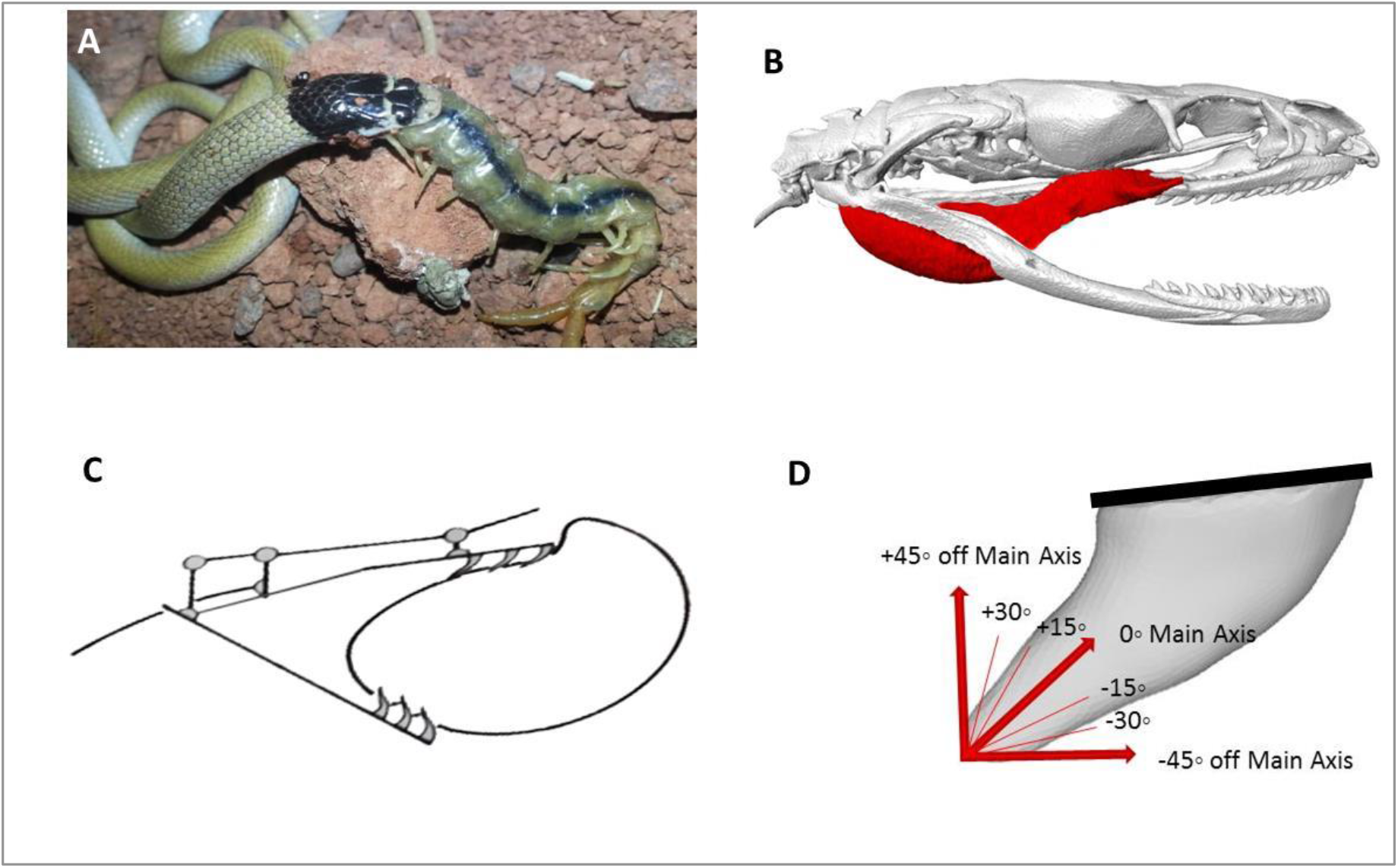
Feeding of *Eirenis persicus* on a massive, elongated arthropod; a centipede (photo by S. Samy); Anatomical position of the pterygoideus and pterygoideus accessories muscles that generate the movement of the maxilla (B); schematic presentation of snake ingestion, showing maxillary teeth sliding over the prey to align the forces acting on the teeth with the long axes of the teeth (C); a range of forces acting in the sagittal plane including the main axis of the tooth, as well as ±45° off the main axis, with the intervals of 15° was implemented (D).

### FEA model

The Finite Element Analysis was performed using a Static Structural Module in ANSYS v 15.0 (ANSYS, Canonburg, PA, USA). To compare the structural strength of an *E. persicus* and a *D. schmidti* maxillary tooth of mean shape in response to the stress caused during biting, the von Mises stress distributions was investigated. von Mises stress is widely accepted for identifying the potential location of the failure due to the stress concentrations in biological structures (Dumont et al., 2005; Whitenack et al., 2011). To be able to compare the von Mises values independent of size differences in the teeth of both examined species, the maxillary tooth of *Dolichophis schmidti* was scaled with the volume of *E. persicus* tooth, and then the constraints, forces and loading conditions described above were implement for both of the examined teeth.

### Fatigue analysis

To check the sensitivity of the tooth to force during repeated loading cycles, we also performed a fatigue analysis. Constraints, forces and loading conditions were set up as defined above, along the main axis of the tooth, but the magnitude of the force was changed from 50% of the defined load, up to 150% of the defined load. The fatigue curve of human dentin was generated based on a study by Nalla and co-workers (Nalla et al., 2003). The resulted fatigue sensitivity chart indicates how fatigue results change as a function of the force magnitude implemented at the tip of the tooth. Fatigue analysis was performed using Fatigue Tools Module of ANSYS v 15.0 0 (ANSYS, Canonburg, PA, USA).

## Results

### Shape analysis

Bonferroni-corrected *P*-values resulting from the nonparametric multivariate analysis of variance of the landmark data of the examined teeth revealed that each group of the teeth (maxillary, palatine, pterygoid and dentary) are significantly different among the examined snakes (except the dentary teeth of the adult *D. schmidti* which are similar to the dentary teeth of the other examined specimens) (Table S1 in the supporting information). The PCA scatterplots of the landmark data for the maxilla, palatine, pterygoid and dentary teeth in the examined specimens are presented in Figure 4. Maxilla and palatine teeth of the *E. persicus* are blunt and conical shape and show clear differences from the sharp and elongated maxillary and palatine teeth of *E. punctatolineatus* as well as the adult and juvenile *D. schmidti*. A similar difference can be noted for the pterygoid teeth. In contrast, the dentary teeth are not as different (Figure 4). The scatter plot resulting from PC1 vs. centroid size of the teeth indicates that the observed divergence in PC1 does not simply reflect size differences, but also difference in the shape of the teeth (Figure 4). The pairwise Euclidean distance of the centroid PC scores of the maxilla, palatine, pterygoid and dentary teeth (Figure 5, values in supporting information, Table S2) revealed that the lowest shape divergence exists among the adult and juvenile *Dolichophis* snakes, while the highest shape divergence is observed between the pterygoid teeth of *E. persicus* and *E. punctatolineatus*. In comparison, *Dolichophis* (both adult and juvenile) differs from *E. punctatolineatus* in the shape of the pterygoid, dentary, maxilla and palatine *Dolichophis* (both adult and juvenile) when compared with *E. persicus*, shows the highest shape divergence among the maxilla, pterygoid, palatine and dentary.

**Figure 4.**
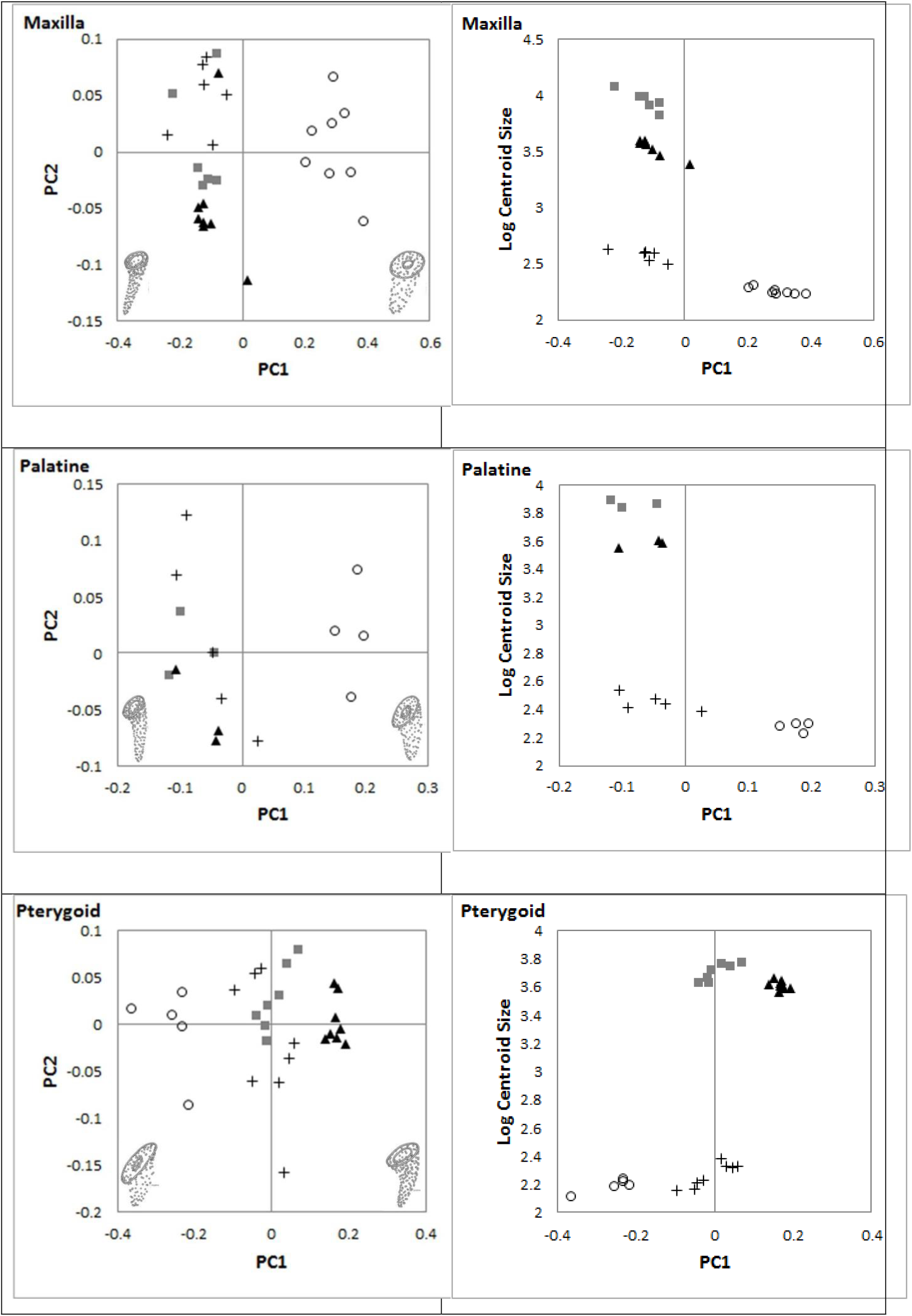
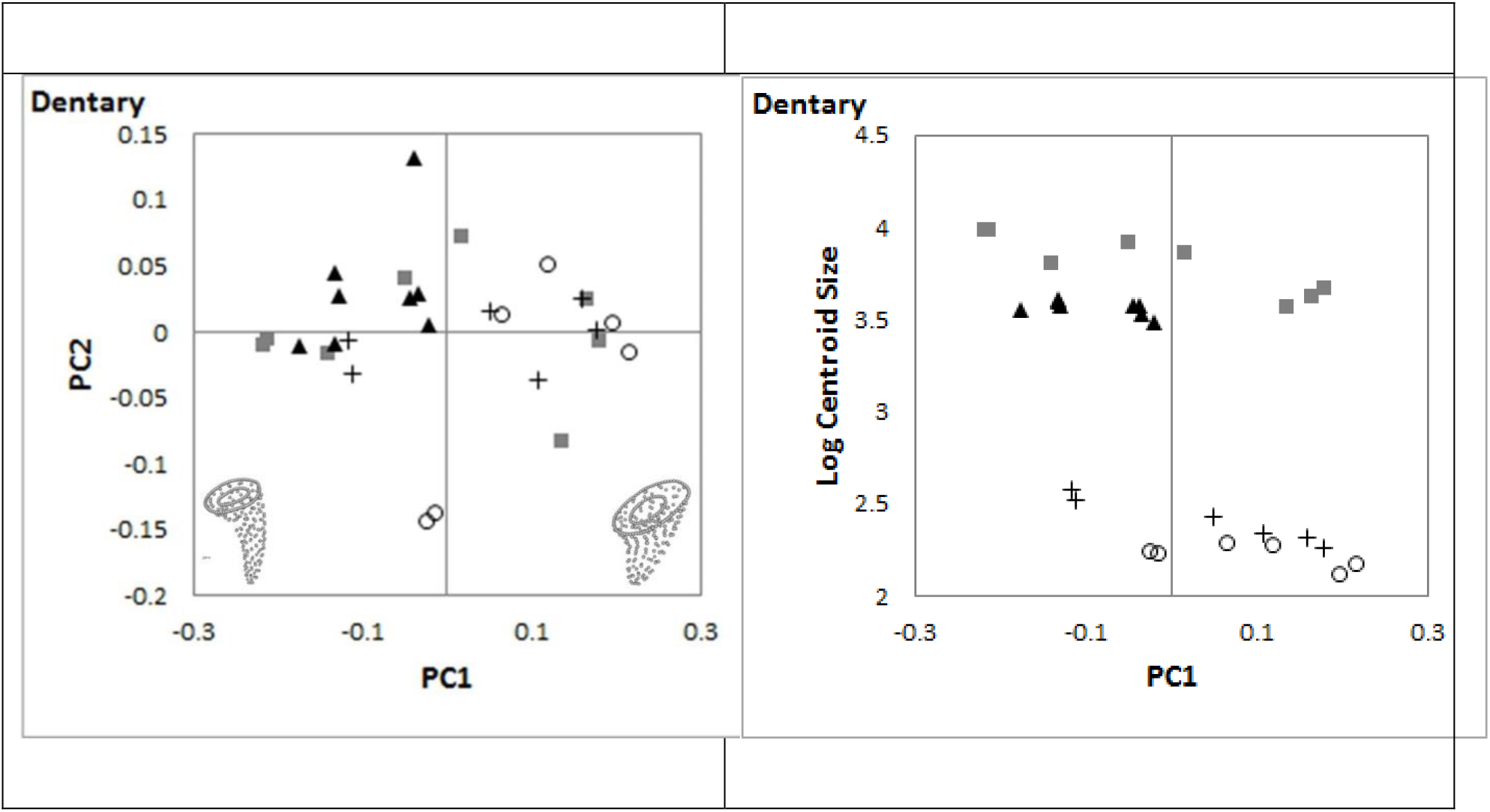
Scatterplot resulting from the principal component analysis on landmark data of teeth of maxilla, palatine, pterygoid and dentary bones in the adult *Dolichophis schmidti* (square), juvenile *D. schmidti* (plus), *Eirenis punctatolineatus* (triangle) and *E. persicus* (circle).

**Figure 5.**
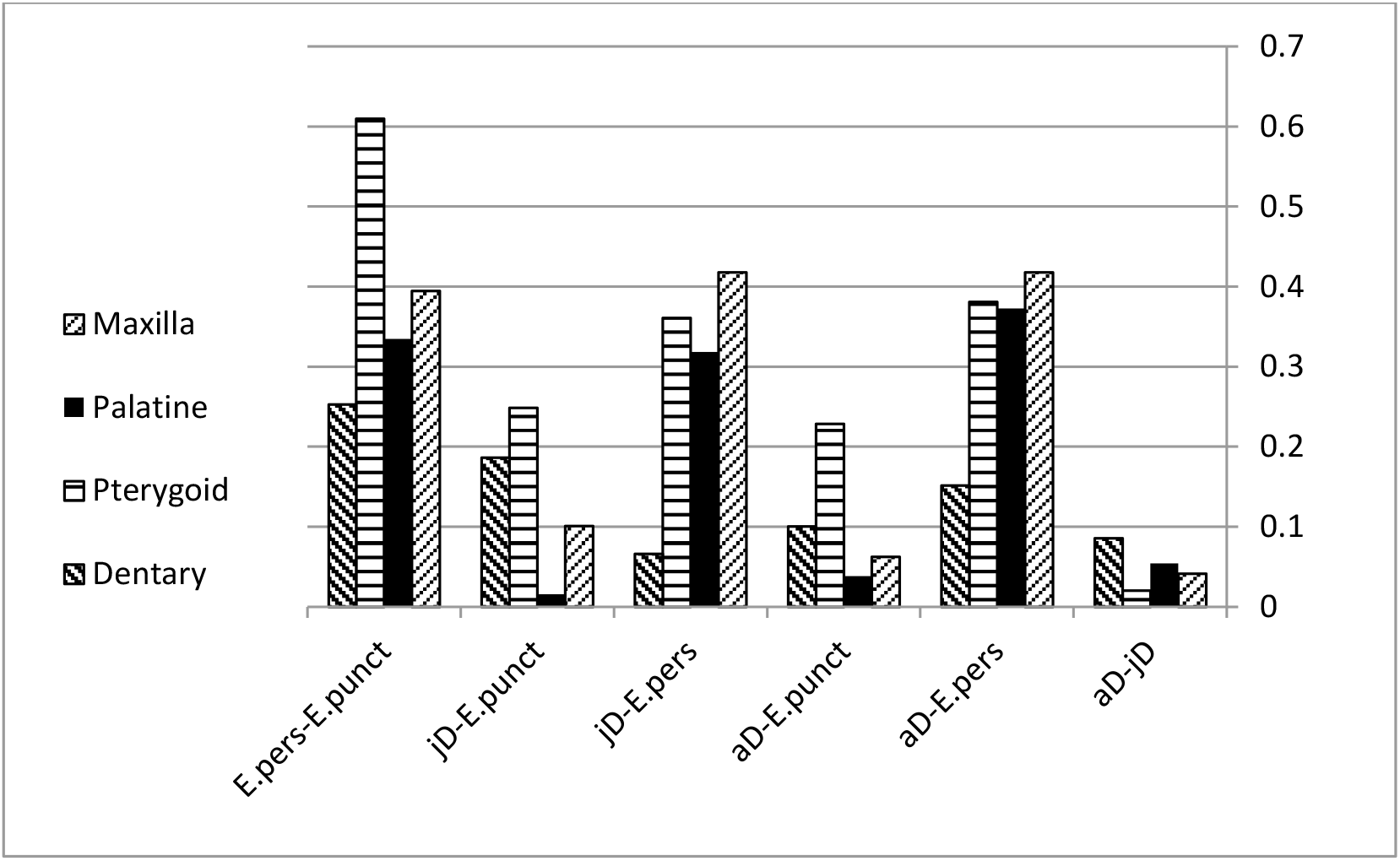
Pairwise Euclidean distance for the maxilla, palatine, pterygoid and dentary teeth in the adult *Dolichophis* (aD), juvenile *Dolichophis* (jD), *Eirenis punctatolineatus* (E.punct) and *Eirenis persicus* (E. pers)

### FEA

When loaded at the tip, both types of teeth regardless of their morphology show the maximum stress at the tip of the teeth (site of loading). While the maximum von Mises stress in the maxillary teeth in *E. persicus* ranges between 41.6 MPa (along −30° of the tooth main axis) to 64.0 MPa (along +45°). For *D. schmidti* the stress ranges between 106.2 MPa (along +15° of the tooth main axis) to 173.2 MPa (along −45°) (Figure 6). In both teeth, the minimum deformation is observed while implementing the force along the main axis of the tooth, while the maximum deformation is observed along the +45° (Figure 7). In both teeth, the stress is more or less restricted to the tip of the teeth while implementing the force along the main axis of the teeth. However, implementing the force along the positive or negative direction leads to confined distribution of stress in the *E. persicus* tooth, but very widely distributed stress in *D. schmidti* tooth (Figure 8).

**Figure 6.**
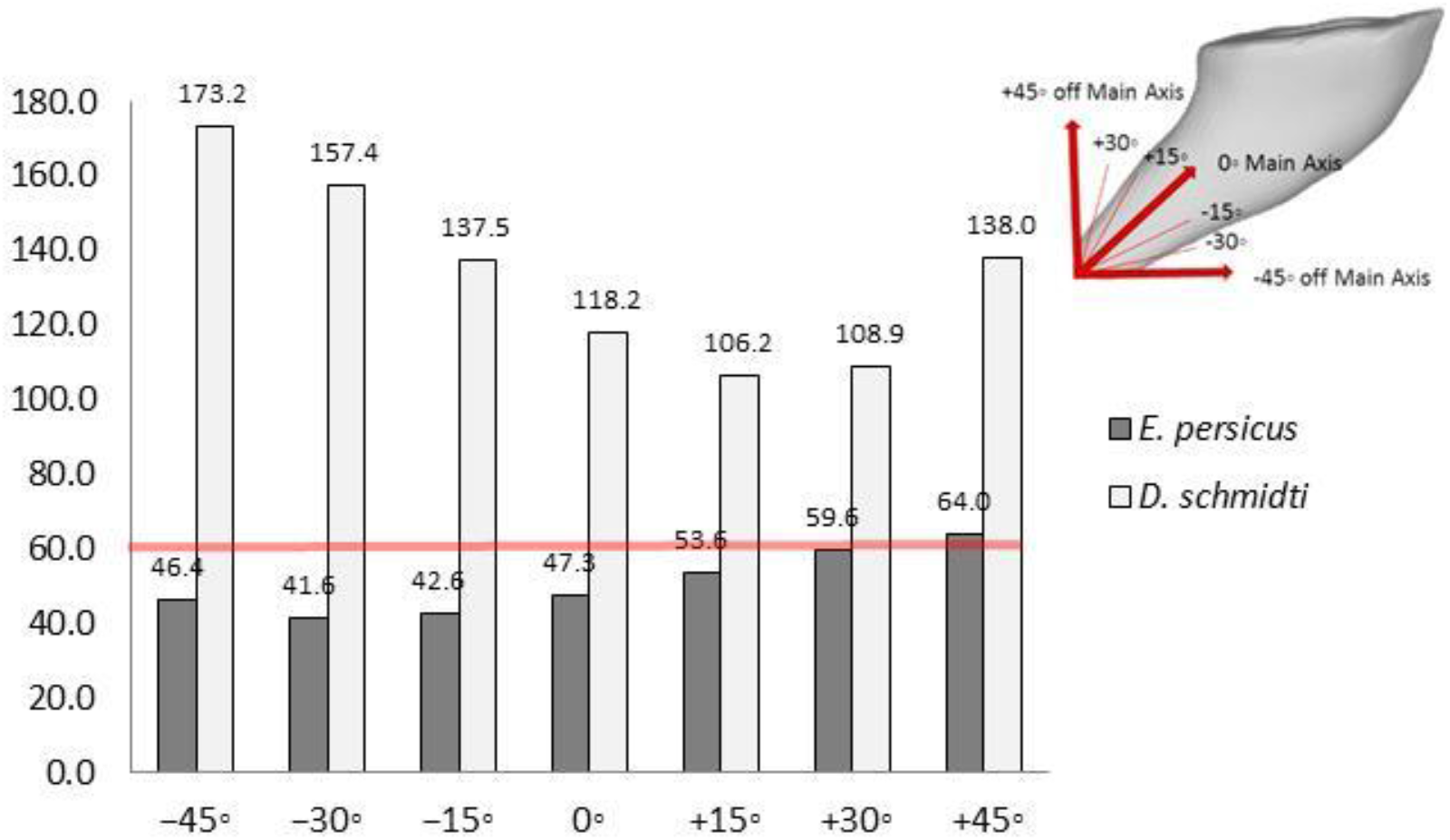
Variation of maximum von Mises stress during loading on tip of the examined mean shape tooth along the main axis of the tooth as well as ±45° off the main axis with intervals of 15°. The red line indicates to von Mises yield criterion of superficial dentin (=61.6) (Giannini et al., 2004).

**Figure 7.**
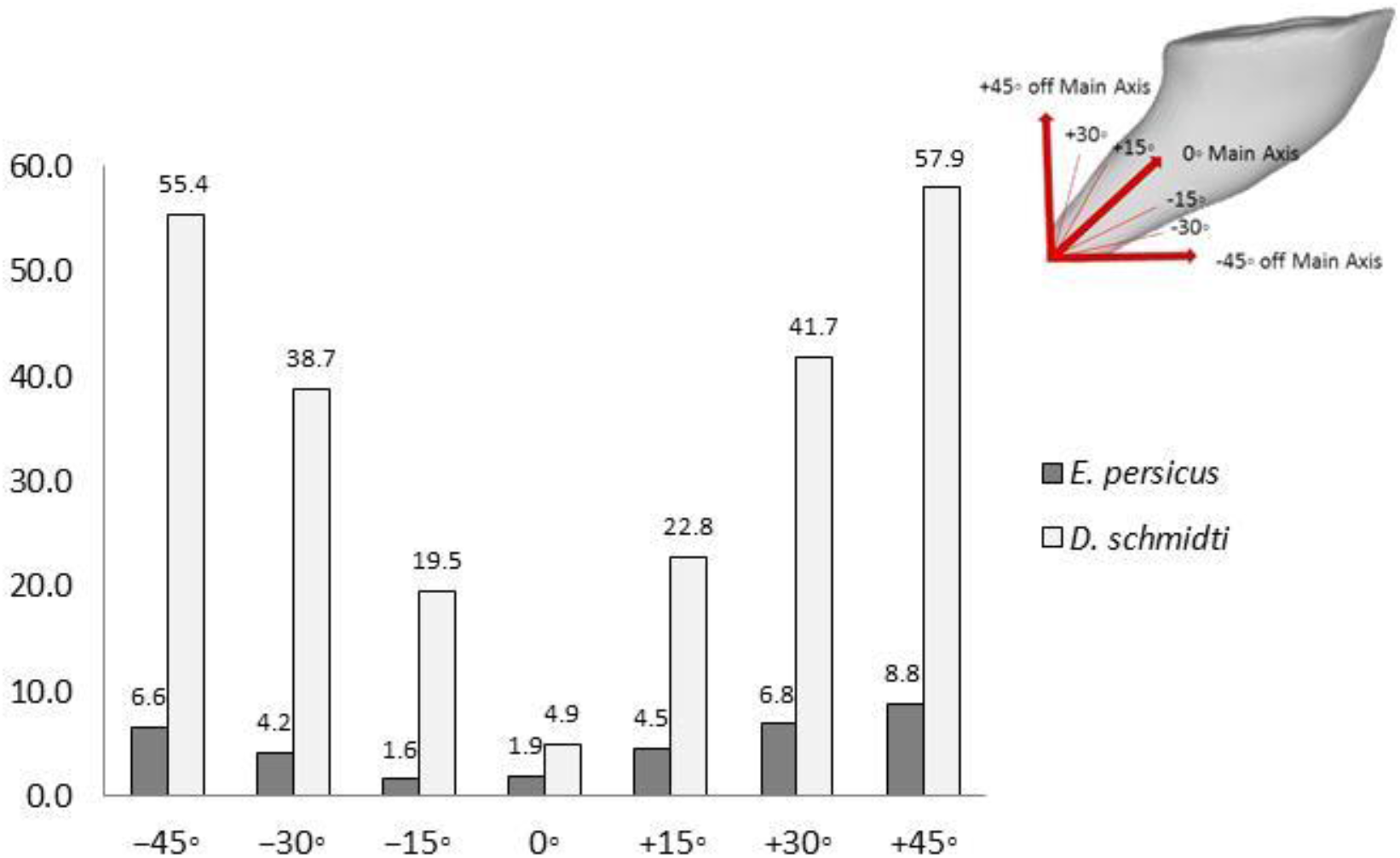
Variation of total deformation (values are in millimeter and has been multiplied in 10^5^) during loading on tip of the examined mean shape tooth along the main axis of the tooth as well as ±45° off the main axis with intervals of 15°.

**Figure 8.**
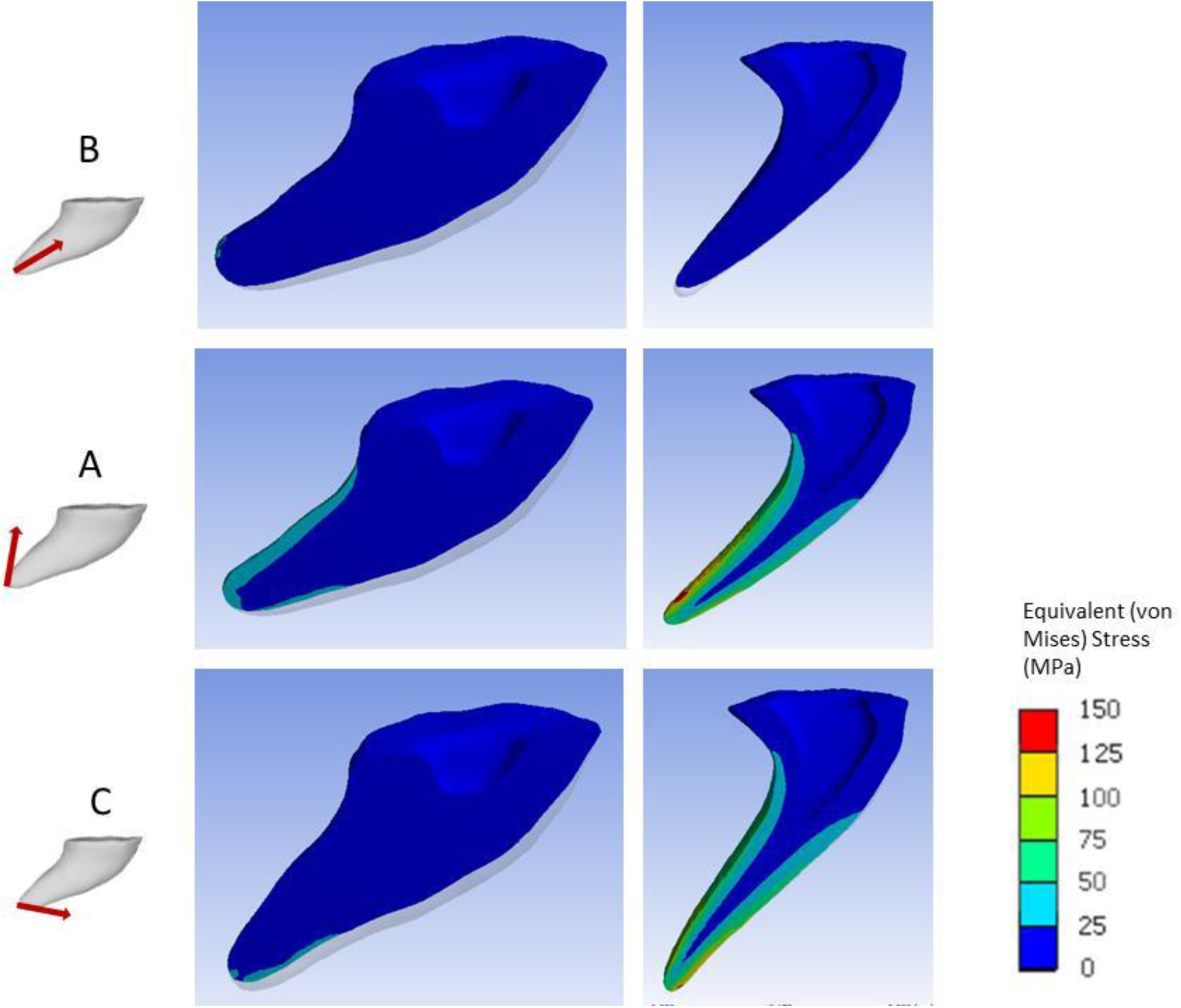
Sagittal surface of *E. persicus* (left) and *D. schmidti* (right) mean shape maxillary teeth, showing the stress distributions (von Mises) resulting from the force loaded to the tip of the tooth along the mains axis (A) of the tooth, +45° (B) and −45° (C) off the main axis.

The fatigue sensitivity chart (Figure 9) indicates that the lifecycle of *E. persicus* teeth is nearly constant when the force magnitude implemented to the tip of the tooth changes between 50% to 150% of the defined force and is 235000 cycles (except for 150% that is 233970). However, for *D. schmidti* the tooth is very sensitive to the defined force magnitude and when the force exceeds 60% of the defined force, the life cycle of the tooth drastically reduces. Hence, at the point of the defined force, the lifecycle of the tooth reduces to around 4883 and around 120% of the defined force, the lifecycle of the tooth falls to zero.

**Figure 9.**
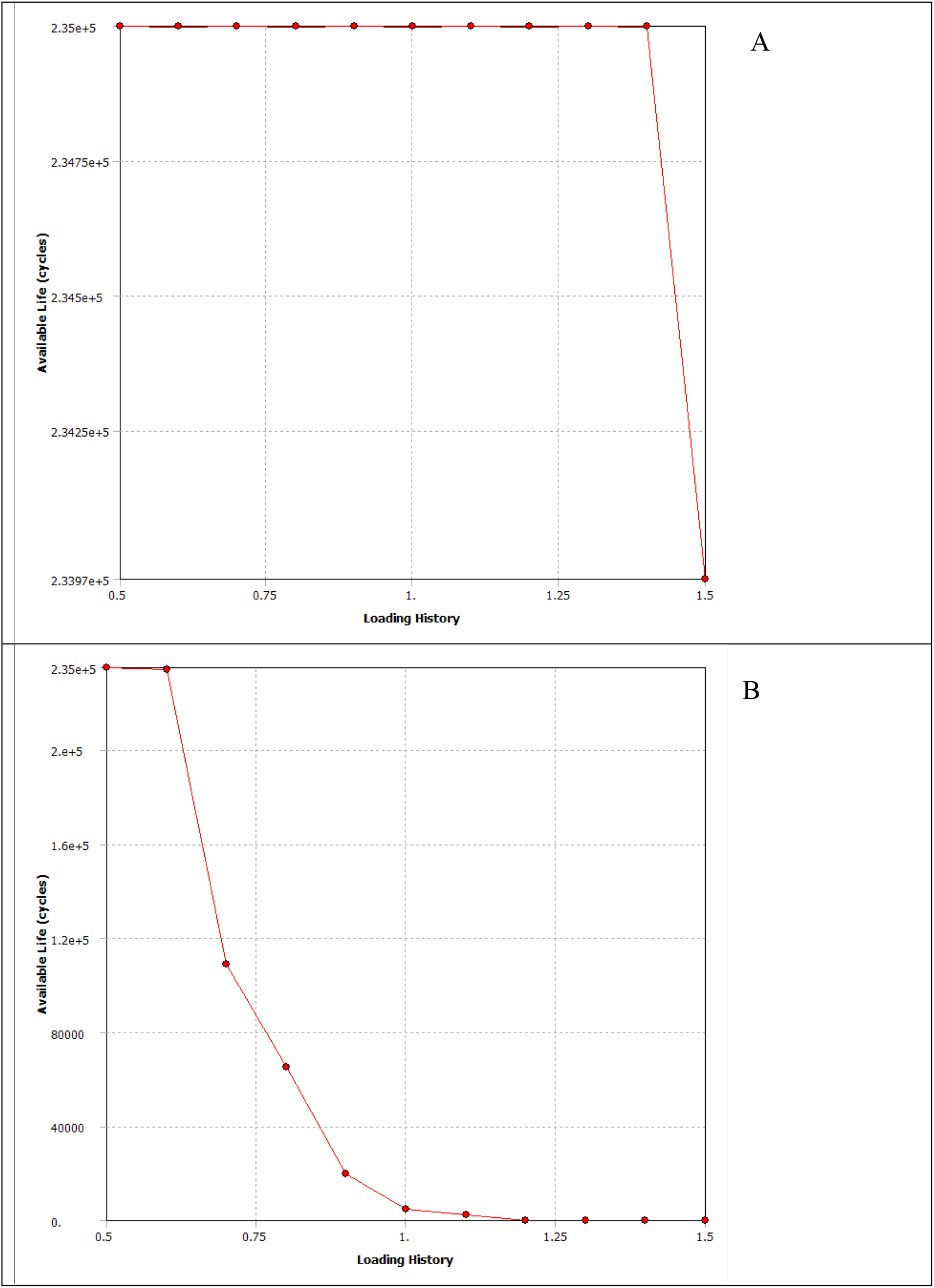
Fatigue sensitivity chart of *E. persicus* (A) and *D. schmidti* (B) when implementing the force along the main axis of the tooth.

## Discussion

The morphology of an organism is controlled by the interaction between historical factors and selective pressures (Losos and Miles, 1994). Variation in the shape observed in the dentary and palatomaxillary teeth of *Dolichophis* and *Eirenis* snakes suggest a possible trade-off between the dietary selective pressures driving adaption of the tooth shape relative to the diet, and the ancestral tooth shape (Knox and Jackson, 2010). Consequently, the adaptive nature of the dentary and palatomaxillary teeth in the examined snakes can only be evaluated based on analyses of their function during feeding.

The first step in snake feeding is the ingestion of the prey which is performed mainly by maxillary teeth. Contraction of pterygoidous muscle retract the maxilla coupled to an elevation of the mandible via the external and internal adductor muscles (Cundall, 1983; Schwenk, 2000). Sharp and elongated maxillary teeth, as observed in *Dolichophis* and *E. punctatolineatus*, are specialized for puncturing or lacerating the skin of the prey, causing successive bleeding or mediate the penetration of secretions of the oral glands inside the prey’s body (Kardong, 1979; Kardong, 1980; Kroll, 1976; Platt, 1967). These kind of teeth are also observed in piscivorous snakes (Savitzky, 1983), and also, the specialized fangs of proteroglyphous, solenoglyphous and opisthoglyphous snakes (Berkovitz and Shellis, 2016).

Shortened, blunt and conically shaped maxillary teeth, as observed in *E. persicus*, seem to be more adapted for biting hard bodied, arthropod prey. Highly shortened (vestigial) and blunt teeth are observed in egg-eating snakes (Gans, 1952). In addition, blunt, molariform dentition, associated with durophagy, has been observed in molluscivorous lizards *e.g.* some amphisbaenians (Pregill, 1984); *Chamaeleolis* lizards (Herrel and Holanova, 2008), northern caiman lizard (*Dracaena guianensis*) (Dalrymple, 1979) and Nile monitors (*Varanus niloticus*) (Rieppel and Labhardt, 1979). Despite some discussion about whether feeding on arthropods should be regarded as durophagy (Savitzky, 1983), the mechanical properties of the exoskeleton of some prey eaten by *Eirenis* snakes are likely putting constraints on tooth shape

The results from our biomechanical modeling confirm that during biting the generated stresses in *E. persicus* are mostly confined to the tip of the tooth and mostly well below the von Mises yield criterion of human dentine that is equal to 61.6 MPa (Giannini et al., 2004). Moreover, our fatigue analysis indicates that *E. persicus* teeth have a high life cycle duration and likely do not break during biting. In contrary, *D. schmidti* teeth appear less well suited for biting a hard prey since the generated stresses widely distribute across the tooth with values roughly 2 to 3 times higher than the von Mises yield criterion of the dentine if its teeth would be subjected to exerting the same forces as those of *E. persicus*. The fatigue analysis further indicates that teeth in *D. schmidti* have a very low life cycle duration when biting and may actually break when loaded similar to those of *E. persicus*.

During the ingestion, the mandible elevates to keep the prey pressed against the month roof, including the upper jaw that is responsible for the main prey transport (Cundall, 1983; Cundall and Deufel, 1999; Schwenk, 2000). A lower degree of specialization was observed among the dentary teeth in the examined snakes suggesting a similar functional constraint in pushing the prey against the upper tooth rows while preventing escape.

In colubroids, swallowing the prey is performed via intraoral transport driven by the fore-aft movement of the medial upper jaws (palatine and pterygoid), coupled with the action of the maxilla and lower jaw (Schwenk, 2000). Intraoral transport implies a coordinated advancing of the jaw over the prey through the action of the protractor and levator muscles on one side, and then clamping down on the prey followed by a retraction of the palate-maxillary unit. Next the process is repeated on the other side resulting in the characteristic pterygoid walk of snakes. Teeth are important to lock the medial upper jaw teeth onto the prey surface (Cundall, 1983). Similar to maxillary teeth, palatine and pterygoid teeth are also short, blunt and conically shape in *E. persicus*, possibly an adaptation for biting and clamping down on arthropod exoskeletons without breaking. Whereas the short and blunt palatopterygoid teeth of *E. persicus* resist loading well, the elongated and sharp teeth in *E. punctatolineatus* and *Dolichophis* are likely more efficient in penetrating the skin of small vertebrate prey.

The toughness of foods plays a crucial role in shaping teeth (Lucas, 2004). Despite the above mentioned examples in snakes, there are well known dental adaptations in relation to diet hardness in other vertebrates too. Herrel et al. (2004) reported tooth shape difference in omnivorous lacertid lizards than their insectivorous counterparts. Omnivores have wider teeth with a larger number of cusps associated, hence a larger tooth perimeter and surface area when compared with insectivores. Blunt molariform teeth are also observed in snail-crushing teiid lizard genus *Dracaena*, snail-eating amphisbaenid lizard and chameleon that are important to avoid tooth breakage while handling a snail (Herrel and Holanova, 2008). Finite element analyses on shark teeth (Whitenack et al., 2011) and spider fang (Bar-On et al., 2014) revealed that teeth loaded in puncture, localized the stress concentrations at the cusp apex. In unicuspid tooth, smoothed tips of the tooth (as observed in blunt teeth of *E. persicus*) reduce the likelihood of chipped teeth (Lawn et al., 2013). Increased tip surface of the tooth via larger number of short, tick, cusps reduce the likelihood of tooth tip failure while loading with a hard prey (Crofts, 2015). Simulations on spider fang clearly showed that conic shape of the tooth, instead of narrow and elongated tooth, is highly adapted structure for stiffness and damage resilience while biting, that limit the stress at the tooth tip and diminish it rapidly away from the apex (Bar-On et al., 2014).

In conclusion, teeth in Aglyphous snakes show adaptation in relation to prey hardness. In Aglyphous snakes feeding solely on arthropods, e.g. *Eirenis persicus*, occurrence of conic shape and blunt maxillary, palatine and pterygoid teeth is a kind of dentary adaptation to well resist the tension of biting a prey, having hard exoskeleton. This adaptation is in contrary to occurrence of elongated and sharp palate-maxillary teeth in counterpart snakes feeding on endotherms or a mix of ectotherms and endotherms, e.g. *Eirenis punctatolineatus* and *Dolichophis schmidti* (adult and juvenile) which is better suited for puncturing or lacerating the skin of the prey.

## Acknowledgements

We are grateful to Dominique Adriaens in Ghent University for his kind collaboration. MR would like to thank the French Embassy in Tehran for a visiting scholar grant (2018) that allowed him to work at the MNHN in Paris, as well as the Naturalis Biodiversity Center, Leiden, The Netherlands for a Martin Fellowship grant.

## Supporting Information

**Table S1.**
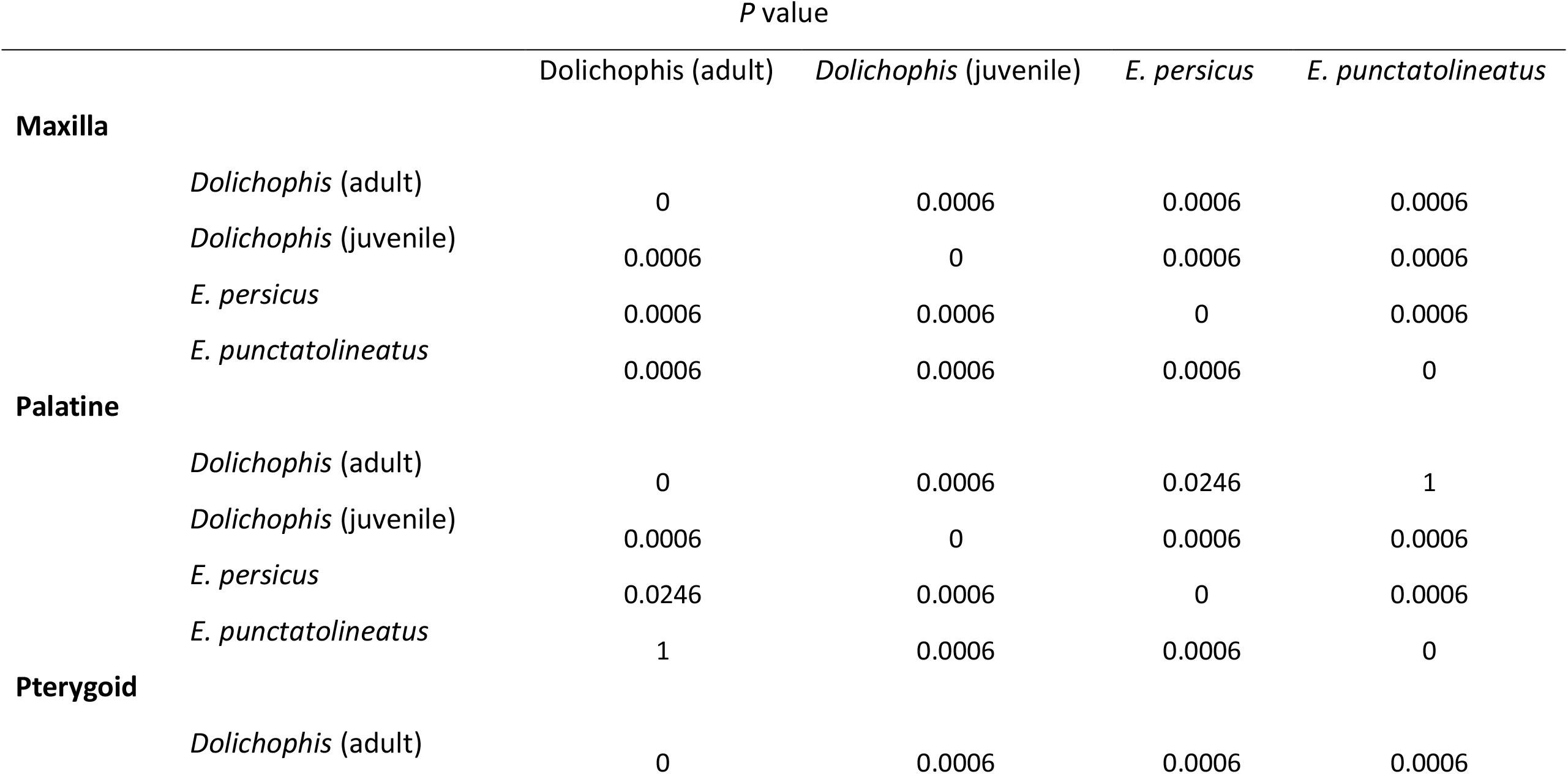
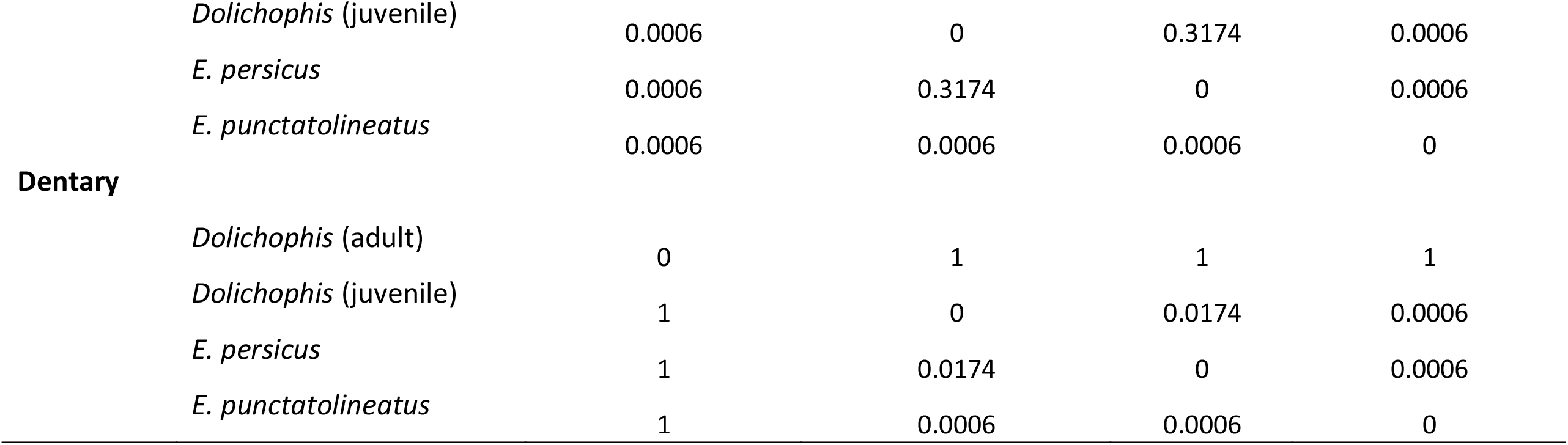
Bonferroni-corrected *P*-values resulting from the nonparametric multivariate analysis of variance on the landmark data of maxilla, palatine, pterygoid and mandible teeth in the examined snakes. Significant values are marked by *.

**Table S2.** Pairwise Euclidean distance along the centroid PC scores of the maxilla, palatine, pterygoid and dentary teeth

|  | Dolichophis (adult) | *Dolichophis* (juvenile) | *E. persicus* | *E. punctatolineatus* |
| --- | --- | --- | --- | --- |
| **Maxilla** |  |  |  |  |
| *Dolichophis* (adult) | 0.000 | 0.042 | 0.418 | 0.063 |
| *Dolichophis* (juvenile) |  | 0.000 | 0.418 | 0.101 |
| *E. persicus* |  |  | 0.000 | 0.394 |
| *E. punctatolineatus* |  |  |  | 0.000 |
| **Palatine** |  |  |  |  |
| *Dolichophis* (adult) | 0.000 | 0.054 | 0.373 | 0.038 |
| *Dolichophis* (juvenile) |  | 0.000 | 0.319 | 0.016 |
| *E. persicus* |  |  | 0.000 | 0.334 |
| *E. punctatolineatus* |  |  |  | 0.000 |
| **Pterygoid** |  |  |  |  |
| *Dolichophis* (adult) | 0.000 | 0.020 | 0.381 | 0.228 |
| *Dolichophis* (juvenile) |  | 0.000 | 0.361 | 0.249 |
| *E. persicus* |  |  | 0.000 | 0.610 |
| *E. punctatolineatus* |  |  |  | 0.000 |
| **Dentary** |  |  |  |  |
| *Dolichophis* (adult) | 0.000 | 0.086 | 0.152 | 0.101 |
| *Dolichophis* (juvenile) |  | 0.000 | 0.066 | 0.186 |
| *E. persicus* |  |  | 0.000 | 0.253 |
| *E. punctatolineatus* |  |  |  | 0.000 |

## References

Bar-On, B., Barth, F. G., Fratzl, P. and Politi, Y. (2014). Multiscale structural gradients enhance the biomechanical functionality of the spider fang. Nature communications 5, 3894.

Berkovitz, B. K. and Shellis, R. P. (2016). The teeth of non-mammalian vertebrates: Academic Press.

Bessone, L. M., Bodereau, E. F., Cabanillas, G. and Dominguez, A. (2014). Analysis of biomechanical behaviour of anterior teeth using two different methods: Finite element method and experimental tests. Engineering 6, 148.

Botton-Divet, L., Cornette, R., Fabre, A.-C., Herrel, A. and Houssaye, A. (2016). Morphological analysis of long bones in semi-aquatic mustelids and their terrestrial relatives. Integrative and comparative biology 56, 1298–1309.

Botton-Divet, L., Houssaye, A., Herrel, A., Fabre, A.-C. and Cornette, R. (2015). Tools for quantitative form description; an evaluation of different software packages for semi-landmark analysis. PeerJ 3, e1417.

Crofts, S. (2015). Finite element modeling of occlusal variation in durophagous tooth systems. Journal of Experimental Biology 218, 2705–2711.

Cundall, D. (1983). Activity of head muscles during feeding by snakes: a comparative study. American Zoologist 23, 383–396.

Cundall, D. and Deufel, A. (1999). Striking patterns in booid snakes. Copeia, 868–883.

Dalrymple, G. (1979). On the jaw mechanism of the snail-crushing lizards: Dracaena.

Deufel, A. and Cundall, D. (2006). Functional plasticity of the venom delivery system in snakes with a focus on the poststrike prey release behavior. Zoologischer Anzeiger-A Journal of Comparative Zoology 245, 249–267.

Dumont, E. R., Piccirillo, J. and Grosse, I. R. (2005). Finite‐element analysis of biting behavior and bone stress in the facial skeletons of bats. The Anatomical Record Part A: Discoveries in Molecular, Cellular, and Evolutionary Biology: An Official Publication of the American Association of Anatomists 283, 319–330.

Gans, C. (1952). The functional morphology of the egg-eating adaptations in the snake genus Dasypeltis. Zoologica 37, 209–244.

Giannini, M., Soares, C. J. and de Carvalho, R. M. (2004). Ultimate tensile strength of tooth structures. Dental Materials 20, 322–329.

Göçmen, B., Werner, Y. L. and Elbeyli, B. (2008). Cannibalism in *Dolichophis jugularis* (Serpentes: Colubridae): More than Random? Current Herpetology 27, 1–7.

Gunz, P. and Mitteroecker, P. (2013). Semilandmarks: a method for quantifying curves and surfaces. Hystrix, the Italian Journal of Mammalogy 24, 103–109.

Gunz, P., Mitteroecker, P. and Bookstein, F. L. (2005). Semilandmarks in three dimensions. In Modern morphometrics in physical anthropology, pp. 73–98: Springer.

Herrel, A. and Holanova, V. (2008). Cranial morphology and bite force in Chamaeleolis lizards–adaptations to molluscivory? Zoology 111, 467–475.

Herrel, A., Vanhooydonck, B. and Van Damme, R. (2004). Omnivory in lacertid lizards: adaptive evolution or constraint? Journal of evolutionary biology 17, 974–984.

Jackson, K., Underwood, G., Arnold, E. and Savitzky, A. H. (1999). Hinged teeth in the enigmatic colubrid, Iguanognathus werneri. Copeia, 815–818.

Kardong, K., Dullemeijer, P. and Fransen, J. (1986). Feeding mechanism in the rattlesnake *Crotalus durissus*. Amphibia-Reptilia 7, 271–302.

Kardong, K. V. (1979). ‘Protovipers’ and the Evolution of Snake Fangs. Evolution, 433–443.

Kardong, K. V. (1980). Evolutionary patterns in advanced snakes. American Zoologist 20, 269–282.

Kardong, K. V. and Young, B. A. (1996). Dentitional surface features in snakes (Reptilia: Serpentes). Amphibia-Reptilia 17, 261–276.

Knox, A. and Jackson, K. (2010). Ecological and phylogenetic influences on maxillary dentition in snakes. Phyllomedusa: Journal of Herpetology 9, 121–131.

Koussoulakou, D. S., Margaritis, L. H. and Koussoulakos, S. L. (2009). A curriculum vitae of teeth: evolution, generation, regeneration. International journal of biological sciences 5, 226.

Kroll, J. C. (1976). Feeding adaptations of hognose snakes. The Southwestern Naturalist, 537–557.

Lawn, B. R., Bush, M. B., Barani, A., Constantino, P. J. and Wroe, S. (2013). Inferring biological evolution from fracture patterns in teeth. Journal of theoretical biology 338, 59–65.

Lelièvre, H., Legagneux, P., Blouin-Demers, G., Bonnet, X. and Lourdais, O. (2012). Trophic niche overlap in two syntopic colubrid snakes (Hierophis viridiflavus and Zamenis longissimus) with contrasted lifestyles. Amphibia-Reptilia 33, 37–44.

Losos, J. B. and Miles, D. B. (1994). Adaptation, constraint, and the comparative method: phylogenetic issues and methods. Ecological morphology: integrative organismal biology 60, 98.

Lucas, P. W. (2004). Dental functional morphology: how teeth work: Cambridge University Press.

Masschaele, B., Cnudde, V., Dierick, M., Jacobs, P., Van Hoorebeke, L. and Vlassenbroeck, J. (2007). UGCT: New X-ray radiography and tomography facility. Nuclear Instruments and Methods in Physics Research Section A: Accelerators, Spectrometers, Detectors and Associated Equipment 580, 266–269.

Nalla, R., Imbeni, V., Kinney, J., Staninec, M., Marshall, S. and Ritchie, R. (2003). In vitro fatigue behavior of human dentin with implications for life prediction. Journal of Biomedical Materials Research Part A: An Official Journal of The Society for Biomaterials, The Japanese Society for Biomaterials, and The Australian Society for Biomaterials and the Korean Society for Biomaterials 66, 10–20.

Platt, D. R. (1967). Natural history of the eastern and the western hognose snakes Heterodon platyrhinos and Heterodon nasicus.

Pregill, G. (1984). Durophagous feeding adaptations in an amphisbaenid. Journal of Herpetology, 186–191.

Rajabizadeh, M. (2018). Snake of Iran. Tehran, Iran: Iranshenasi.

Rajabizadeh, M., Adriaens, D., De Kegel, B., Avci, A., Ilgaz, Ç. and Herrel, A. (2019). Body size miniaturization in a lineage of colubrid snakes: Implications for cranial anatomy. Journal of anatomy In Press.

Rieppel, O. and Labhardt, L. (1979). Mandibular mechanics in Varanus niloticus (Reptilia: Lacertilia). Herpetologica, 158–163.

Savitzky, A. H. (1981). Hinged teeth in snakes: an adaptation for swallowing hard-bodied prey. Science 212, 346–349.

Savitzky, A. H. (1983). Coadapted character complexes among snakes: fossoriality, piscivory, and durophagy. American Zoologist 23, 397–409.

Schlager, S. (2013). Morpho: Calculations and visualisations related to Geometric Morphometrics. R package version 0.23 3.

Schwenk, K. (2000). Feeding: form, function and evolution in tetrapod vertebrates: Academic Press.

Team, R. C. (2018). R Foundation for Statistical Computing; Vienna, Austria: 2014. R: A language and environment for statistical computing, 2013.

Terent év, P. V. and Chernov, S. A. (1965). Key to amphibian and reptiles. Jerusalem, Israel: Program for Scientific translation [English translation of Terent év & Chernov, 1949].

Vlassenbroeck, J., Dierick, M., Masschaele, B., Cnudde, V., Van Hoorebeke, L. and Jacobs, P. (2007). Software tools for quantification of X-ray microtomography at the UGCT. Nuclear Instruments and Methods in Physics Research Section A: Accelerators, Spectrometers, Detectors and Associated Equipment 580, 442–445.

Waters, N. (1980). Some mechanical and physical properties of teeth. The mechanical properties of biological materials, 99–134.

Whitenack, L. B., Simkins Jr, D. C. and Motta, P. J. (2011). Biology meets engineering: the structural mechanics of fossil and extant shark teeth. Journal of morphology 272, 169–179.

Wiley, D. F., Amenta, N., Alcantara, D. A., Ghosh, D., Kil, Y. J., Delson, E., Harcourt-Smith, W., Rohlf, F. J., St John, K. and Hamann, B. (2005). Evolutionary morphing. In VIS 05. IEEE Visualization, 2005., pp. 431–438: IEEE.

Zaher, H. and Rieppel, O. (1999). Tooth implantation and replacement in squamates, with special reference to mosasaur lizards and snakes. Amer. museum novitates.

Zahradnicek, O., Buchtova, M., Dosedelova, H. and Tucker, A. S. (2014). The development of complex tooth shape in reptiles. Frontiers in physiology 5, 74.

Zhang, Y.-R., Du, W., Zhou, X.-D. and Yu, H.-Y. (2014). Review of research on the mechanical properties of the human tooth. International journal of oral science 6, 61.

